# Empagliflozin preserves mitochondrial function and reduces tubular injury in obese type 2 diabetic ZSF-1 rats

**DOI:** 10.64898/2026.03.10.710788

**Authors:** Hannah Weissbach, Meike Seitz, Julia Moosheimer, Florian Gembardt, Antje Schauer, Anita Männel, Michael P. Pieper, Christian Hugo, Volker Adams, Anne Steglich

**Affiliations:** University Hospital Carl Gustav Carus at the Technische Universität Dresden, Department of Internal Medicine III, Division of Nephrology, Dresden, Germany; Heidelberg University, Mannheim Medical Faculty, 5th Department of Medicine, Theodor-Kutzer-Ufer 1-3, 68167 Mannheim, Germany; Laboratory of Molecular and Experimental Cardiology, TU Dresden, Heart Center Dresden, 01307 Dresden, Germany; CardioMetabolic Diseases Research, Boehringer Ingelheim Pharma GmbH & Co KG, Biberach an der Riss, Germany

**Keywords:** diabetic nephropathy, SGLT2, empagliflozin, ZSF-1 rat model, mitochondrial dysfunction

## Abstract

Diabetic nephropathy remains the leading cause of end-stage renal disease. The ZSF-1 rat model combines features known as the metabolic syndrome, such as type 2 diabetes mellitus, hypertension and obesity, developing progressive kidney disease. In this study, we investigated the nephroprotective mechanisms of the SGLT2 inhibitor empagliflozin, focusing on mitochondrial function. Obese ZSF-1 rats were randomized at 24 weeks of age to receive either placebo or empagliflozin for eight weeks, while lean ZSF-1 rats served as healthy controls.

Kidney function, assessed by glomerular filtration rate (GFR), was significantly reduced in obese rats and was not improved by empagliflozin treatment. However, obese animals exhibited increased tubular injury, tubular cast formation, and elevated total and tubular proteinuria, all of which were attenuated by empagliflozin. Mitochondrial function was evaluated in freshly isolated cortical kidney mitochondria by measuring oxygen consumption. Obese ZSF-1 rats showed impaired respiratory capacity and reduced protein expression of oxidative phosphorylation (OXPHOS) complexes II, III, IV, and V, indicating mitochondrial dysfunction. Empagliflozin treatment improved mitochondrial function by enhancing complex I- and IV-linked respiration and restoring the expression of OXPHOS complexes II, III, and IV. In addition, empagliflozin treatment was linked to improved mitochondrial dynamics and modulation of autophagic activity, suggesting enhanced mitochondrial quality control.

Overall, these findings demonstrate that empagliflozin exerts nephroprotective effects primarily at the tubular level in obese ZSF-1 rats. The beneficial effects appear to be mediated through improved mitochondrial function, enhanced mitochondrial integrity, and reduced tubular injury.

## Introduction

Chronic kidney disease (CKD) affects more than 10% of the global population and is particularly common among patients with heart failure [1]. The metabolic syndrome, a cluster of conditions including obesity, hypertension and insulin resistance increases the risk for diabetes mellitus type 2 (T2D) and the development of diabetic nephropathy (DN). It also accelerates the progression of diabetic-hypertensive kidney disease by promoting oxidative stress, inflammation and endothelial dysfunction, further worsening renal outcomes [2].

The Zucker Fatty and Spontaneously Hypertensive (ZSF-1) rat, a cross between Zucker Diabetic Fatty (ZDF) female rats and lean male Spontaneously Hypertensive Heart Failure (SHHF) rats, develops type 2 diabetes mellitus with comorbidities such as hypertension, obesity and heart failure at 10 to 20 weeks of age [3]. These rats develop progressive renal damage over time with proteinuria, glomerulosclerosis and tubulointerstitial fibrosis, which closely resemble the renal changes seen in human DN [4]. Overall, the ZSF-1 rat model is well suited for longitudinal studies of renal disease progression and intervention [4,5]. Sodium-Glucose-Like Transporter 2 (SGLT2) inhibitors (SGLT2i), the first anti-diabetic drugs directly acting on the kidney, demonstrated significant overall renoprotective effects in mice and human [6–8]. In addition to improved renal outcomes, clinical trials have shown that SGLT2i act independently of glycemic control to improve cardiovascular and renal outcomes [9].

In experimental DN, characteristics of nephroprotective effects of SGLT2i are reduction of renal and glomerular hypertrophy and proteinuria, as well as amelioration of histological disease progression in glomerular and also in tubulointerstitial areas [8,10,11]. While the beneficial glomerular effects of gliflozins are often attributed to macula densa–mediated restoration of tubuloglomerular feedback, substantial protective effects within the tubulointerstitium may be due to mechanisms involving the regulation of energy metabolism. Tubular epithelial cells are the first to be affected by hyperglycemic injury [12]. Enhanced glucose reabsorption by SGLT2 increases mitochondrial adenosine triphosphate (ATP) production to meet the high energy demand of tubular transport processes. Increased ATP production accounts for increased levels of reactive oxygen species (ROS), resulting in oxidative stress, tubular cell damage, and the activation of pro-inflammatory and pro-fibrotic pathways [12,13]. SGLT2i results in a significant reduction of glucose supply, leading to metabolic reprogramming [14]. The metabolic reprogramming leads to a shift to increased fatty acid oxidation and a subsequent increase in ketone body production [15]. Ketones can be more easily and efficiently utilized by mitochondria, which ameliorates mitochondrial dysfunction and attenuates oxidative stress in kidney [15,16]. To investigate the impact of SGLT2i on mitochondrial dysfunction and kidney disease progression, we measured oxygen consumption rates in freshly isolated renal mitochondria from obese ZSF1 rats to assess oxidative phosphorylation (OXPHOS) capacity. In addition, we evaluated the effect on empagliflozin on tubular reabsorption and health.

## Results

### Basic characteristics of obese ZSF-1 rats

At the age of 24 weeks, the obese ZSF-1 rats were randomized into a placebo-treated and an empagliflozin treated group (Figure 1A). At 32 weeks, obese animals had significantly higher bodyweight compared to lean. Treatment with empagliflozin significantly prevented body weight gain and reduced blood glucose levels. Empagliflozin did not lead to a further enlargement of kidney mass, as renal hypertrophy was already significantly increased in placebo obese ZSF-1 rats (Table 1).

**Figure 1.**
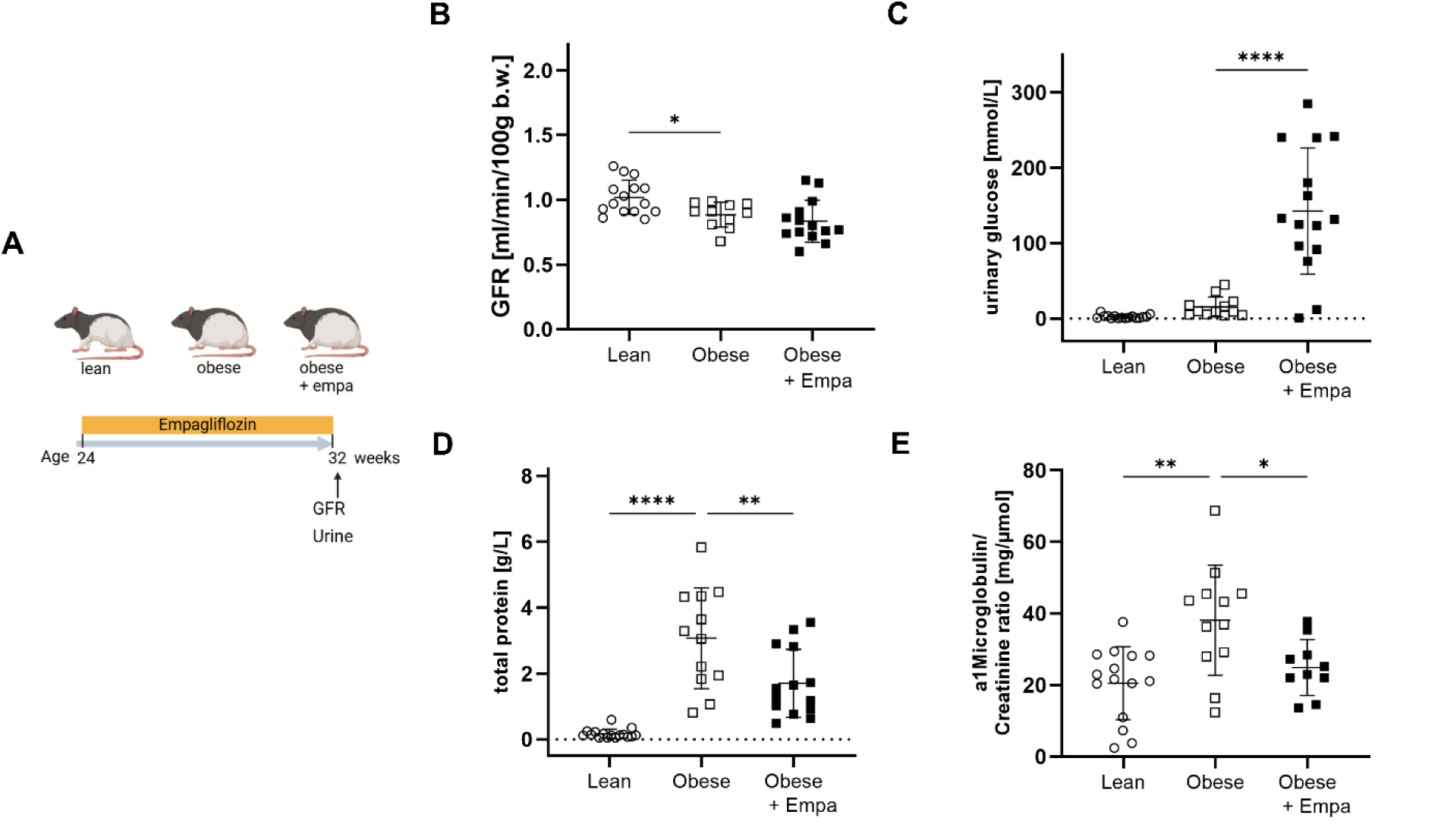
Renal parameters. A) Experimental timeline. Lean and obese ZSF-1 rats were used at the age of 24 weeks. Obese rats were treated with vehicle or empagliflozin for 8 weeks. Lean rats served as control. B) Renal function was assessed in lean (white circles, n = 15), obese (white squares, n = 11) and obese empagliflozin-treated (black squares, n = 14) ZSF-1 rats by measurement of glomerular filtration rate (GFR). Urine was collected to determine C) glucose excretion, D) total protein, E) a1-microglobulin to creatinine ratio in lean (n= 15), obese (n = 12) and empagliflozin-treated obese (n = 13) animals.

**Table 1.** Basic animal characteristics of the 32-week-old lean, obese and obese rats treated with empagliflozin (Obese + Empa. ). KW/Tibia - kidney weight to tibia length ratio.

| Parameter | n | Lean | Obese | Obese + Empa |
| --- | --- | --- | --- | --- |
| Bodyweight [g] | 15 | 255.9 ± 13.7 | 529.6 ± 30.4*** | 491.8 ± 24.4### |
| Blood glucose [mmol/L] | 12 | 18.6 ± 7.0 | 38.2 ± 5.2*** | 25.5 ± 2.5### |
| KW/Tibia [mg/mm] | 14 | 28.2 ± 2.0 | 47.0 ± 4.7*** | 49.4 ± 3.8 |

\*\*\*p < 0.001 vs lean, ###p < 0.001 vs obese

### Renal evaluation of ZSF-1 obese rats regarding diabetic nephropathy

Renal function was evaluated by measuring the glomerular filtration rate (GFR). At 24 weeks of age, GFR measurements in lean and obese rats showed no differences (Supplement Figure S1). However, by 32 weeks, obese rats exhibited a significant reduction in GFR compared with lean controls, which was not restored by empagliflozin treatment (Figure 1B). Urinalysis revealed slightly increased urinary glucose excretion in obese rats (Figure 1C), with pronounced glucosuria following empagliflozin treatment (Figure 1C). Total urinary protein excretion was significantly elevated in obese rats (Figure 1D), whereas empagliflozin treatment significantly improved proteinuria. To further characterise proteinuria, urinary α1-microglobulin excretion was measured. α1-microglobulin excretion was significantly increased in obese rats and was significantly reduced to the level of lean animals after eight weeks of empagliflozin treatment (Figure 1F).

Renal SGLT1 and SGLT2 mRNA expression were also assessed. SGLT1 expression did not differ between groups (Supplement Figure S2A). In contrast, SGLT2 expression was significantly upregulated in obese rats but was not significantly increased anymore after empagliflozin treatment (Supplement Figure S2B).

Histological evaluation of PAS-stained sections demonstrated significant glomerular hypertrophy in obese rats (Figure 2A). Empagliflozin treatment did not attenuate glomerular enlargement. The mean glomerular PAS-positive area did not differ between the groups (Figure 2B). In addition, tubular injury was assessed in PAS-stained sections. Lean control rats exhibited minimal tubular dilation (Figure 2C). In contrast, obese rats showed significantly greater tubular damage, including tubular dilation, loss of brush border, and protein cast formation, which was significantly reduced by empagliflozin treatment (Figure 2D). Quantitative analysis revealed a significantly higher number of protein casts in obese rats in both cortical (Figure 2D) and medullary regions (Figure 2E), which were significantly reduced by empagliflozin treatment. Collagen I and III deposition was evaluated using picrosirius red staining. Obese rats showed similar collagen I and III accumulation compared with lean controls (Figure 2F). Unexpectedly, empagliflozin treatment further increased collagen I and III deposition compared with placebo-treated obese rats. Furthermore, collagen IV was significantly increased in obese rats (Figure 3A). Empagliflozin treatment slowed thickening of the glomerular basement membrane, showing no significant differences to lean or obese rats (Figure 3A).

**Figure 2.**
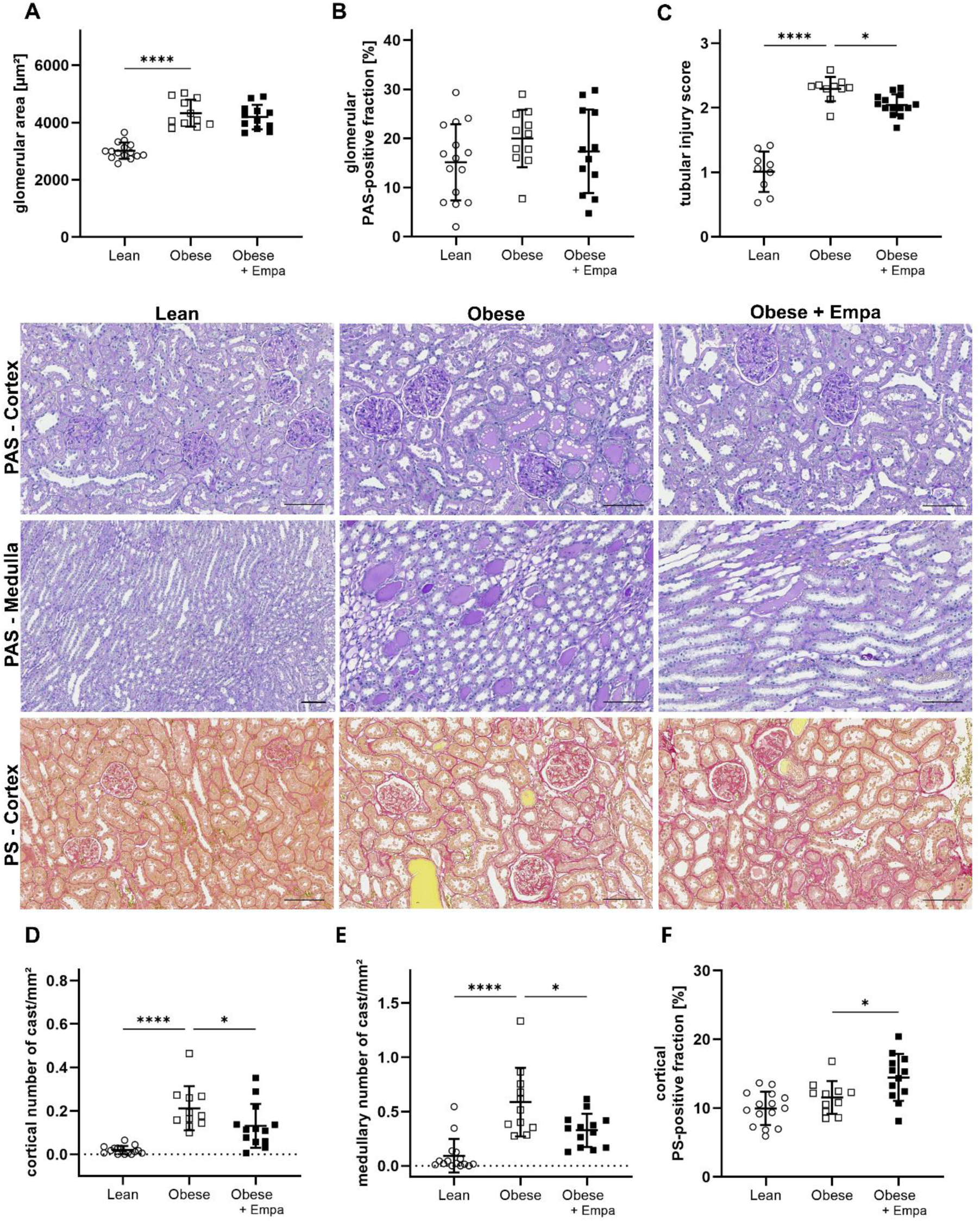
Morphological evaluation of the kidney. Representative images show kidney sections stained with periodic acid–Schiff (PAS) and Picrosirius red (PS). Scale bar represents 100 µm. Morphological evaluation was performed in lean (white circles, n = 15), obese (white squares, n = 11), and empagliflozin-treated obese (black squares, n = 12) ZSF-1 rats. A) Glomerular size expressed as mean area in µm². B) Percentage of PAS-positive glomerular area. C) Tubular injury score. D) Number of cortical protein casts normalized to cortical area. E) Number of medullary protein casts normalized to medullary area. F) Cortical PS-positive fraction.

**Figure 3.**
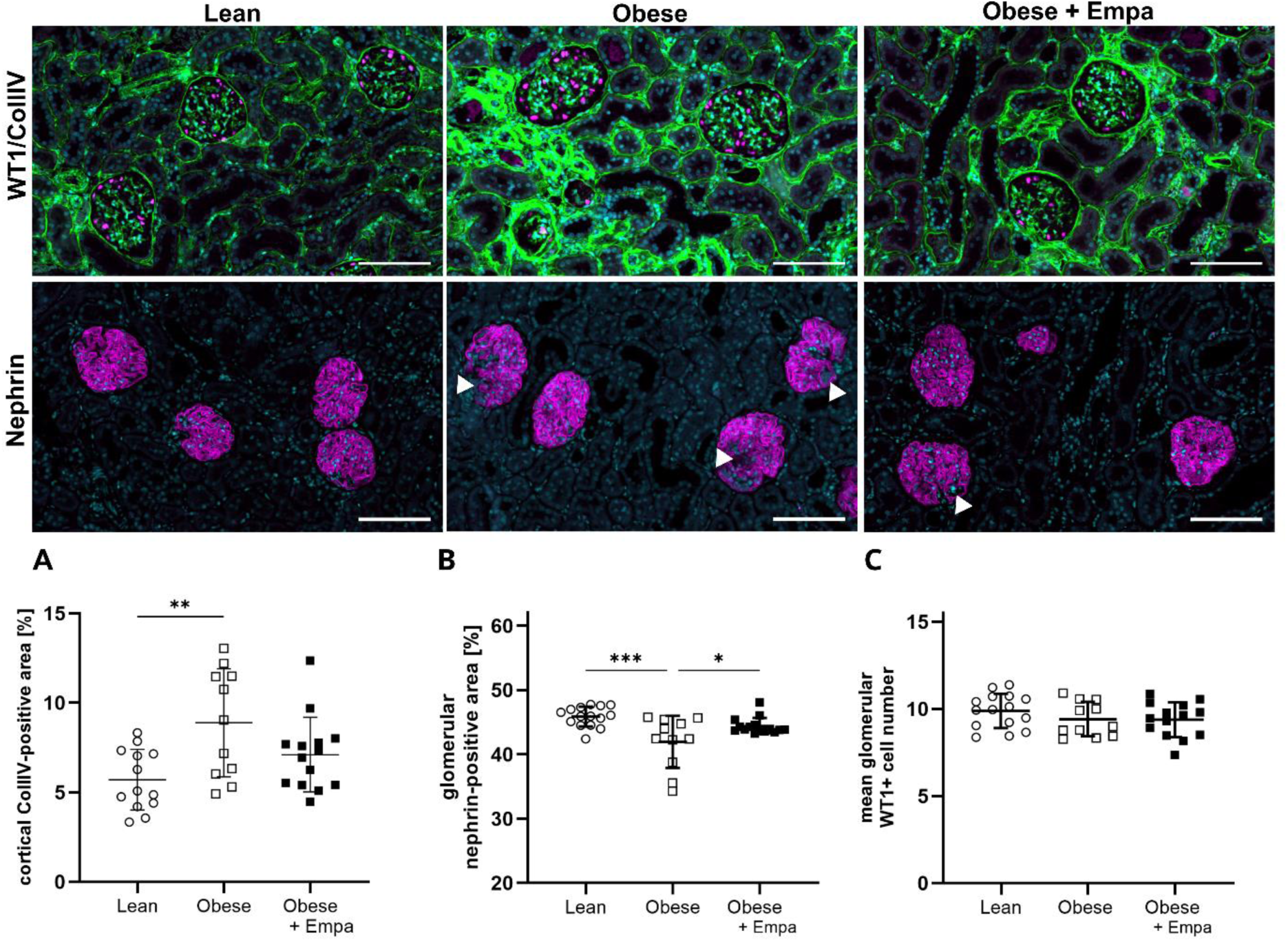
Podocyte dysfunction in ZSF-1 rats. Representative images show kidney sections stained with WT1 (magenta, upper panel) and Collagen IV (green, upper panel) and nephrin (magenta, lower panel). White arrows indicate areas of missing nephrin expression. Scale bar represents 100 µm. A) Quantification of cortical Collagen IV-positive and B) Quantification of glomerular nephrin-positive area and C) Quantification of mean glomerular WT1-positive cell number. Lean (white circles, n = 15, except CollIV n=13), obese (whie squares, n = 11), and empagliflozin-treated obese (black squares, n = 14) ZSF-1 rats.

In addition, podocyte health was evaluated. Nephrin positive area was significantly reduced in obese compared to lean rats (Figure 3B), which was significantly improved by empagliflozin treatment. While nephrin positive area changed, the number of WT1-positive podocytes was unchanged indicating podocyte dysfuction rather than podocyte loss (Figure 3B).

### Mitochondrial function

Mitochondrial function was assessed by oxygen consumption. Stimulation of basal respiration (complex I - state 2 respiration) was not between the groups (Figure 4A). The oxidative phosphorylation after ADP addition (complex I – state 3) was not different in obese rats, but empagliflozin treatment resulted in significantly higher oxygen consumption due to higher ATP production (Figure 4B). Calculating the respiratory control ratio (RCR) for complex I, a significantly higher ratio was observed for the empagliflozin treated obese rats compared to obese rats (Figure 4C).

**Figure 4.**
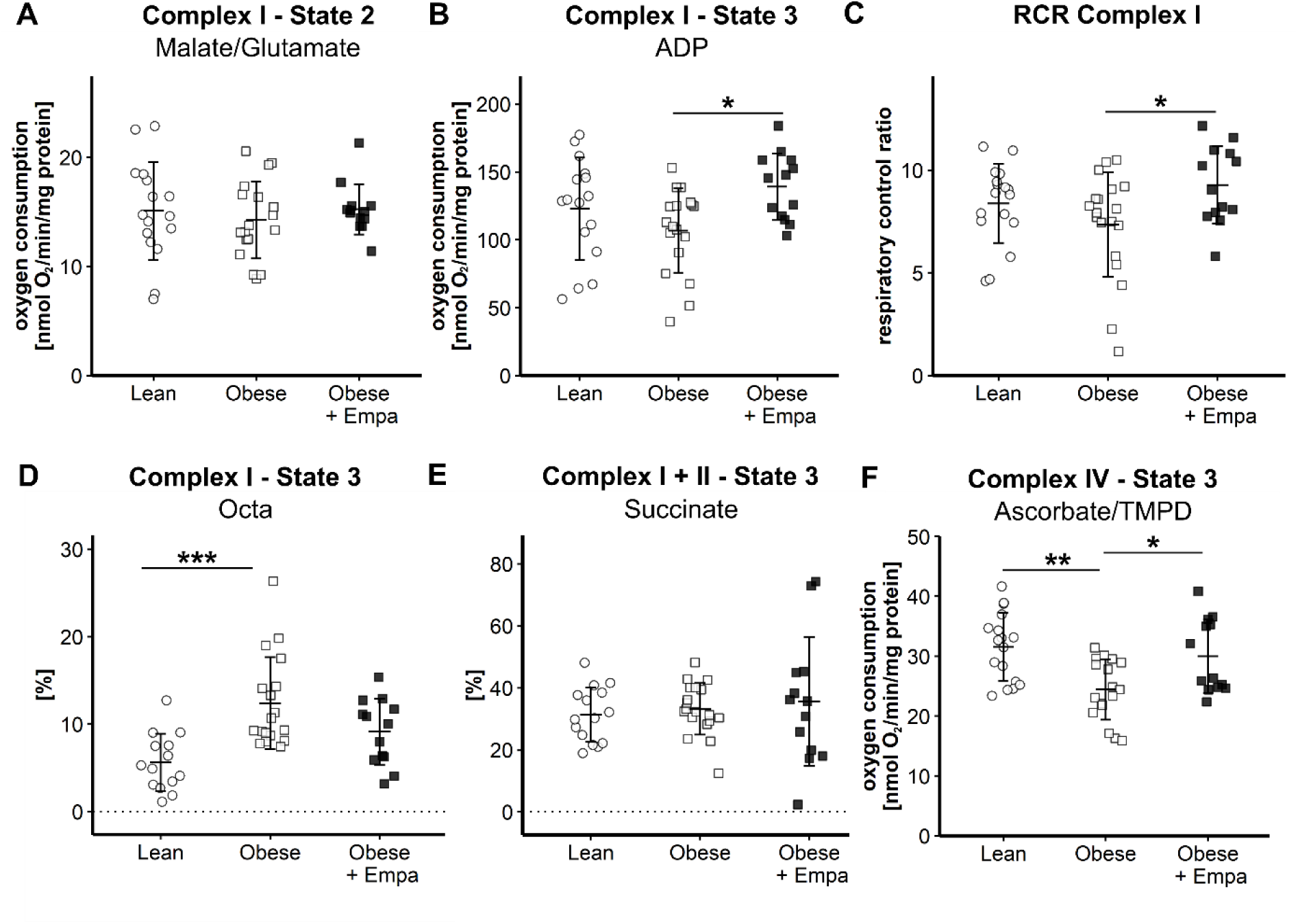
Mitochondrial respiration of complex I, II and IV. Mitochondrial function was assessed in freshly isolated kidney mitochondria of lean (white circles, n = 16), obese (white squares, n = 18) and empagliflozin-treated obese (black squares, n = 14) ZSF-1 rats. A) To measure complex I state 2 leak respiration the substrates malate and glutamate were added. B) By addition of ADP complex I - state 3 respiration was measured. C) The respiratory control ratio (RCR) of complex I was calculated as state 3/state 2 respiration. To measure mitochondrial substrate preferences D) octanoylcarnithine (Octa) for complex I -state 3 respiration and E) succinate for complex I + II respiration state 3 were added. F) Maximal respiration of complex IV was measured after addition of ascorbate and tetramethyl-p-phenylene diamine (TMPD) and inhibition of complex IV by sodium azide.

Octanoylcarnithine supplementation for substrate preference evaluation significantly increased oxygen consumption in the obese compared to lean controls (Figure 4D), and was normalized by empagliflozin treatment. Maximal stimulation of ADP phosphorylation (complex I and II) did not show alterations of oxygen consumption between the groups (Figure 4E). The most present effect for mitochondrial respiration was observed for complex IV. The obese rats had a significant lower oxygen consumption (Figure 4F). The observed effect was diminished by empagliflozin treatment (Figure 4F).

Next, we investigated if the changes in mitochondrial respiration are related to changes in protein expression of the mitochondrial respiratory complexes I-V. Complex I protein expression was unchanged in obese rats compared to lean controls and not influenced by empagliflozin treatment (Figure 5A). The phenotype of the obese rats led to a significantly reduced protein expression of complex II, III and IV (Figure 5B, C, D). Empagliflozin treatment restored protein expression of complex II, III and IV over the treatment period and normalized it to lean level in regard to complex II and III. For complex V, a significant reduction in protein was seen in the obese rats which was normalized by empagliflozin treatment (Figure 5E). Additionally, the cardiolipin content was determined in kidney mitochondria. After empagliflozin treatment the cardiolipin content significantly increased compared to the placebo-treated obese rats (Figure 5F).

**Figure 5.**
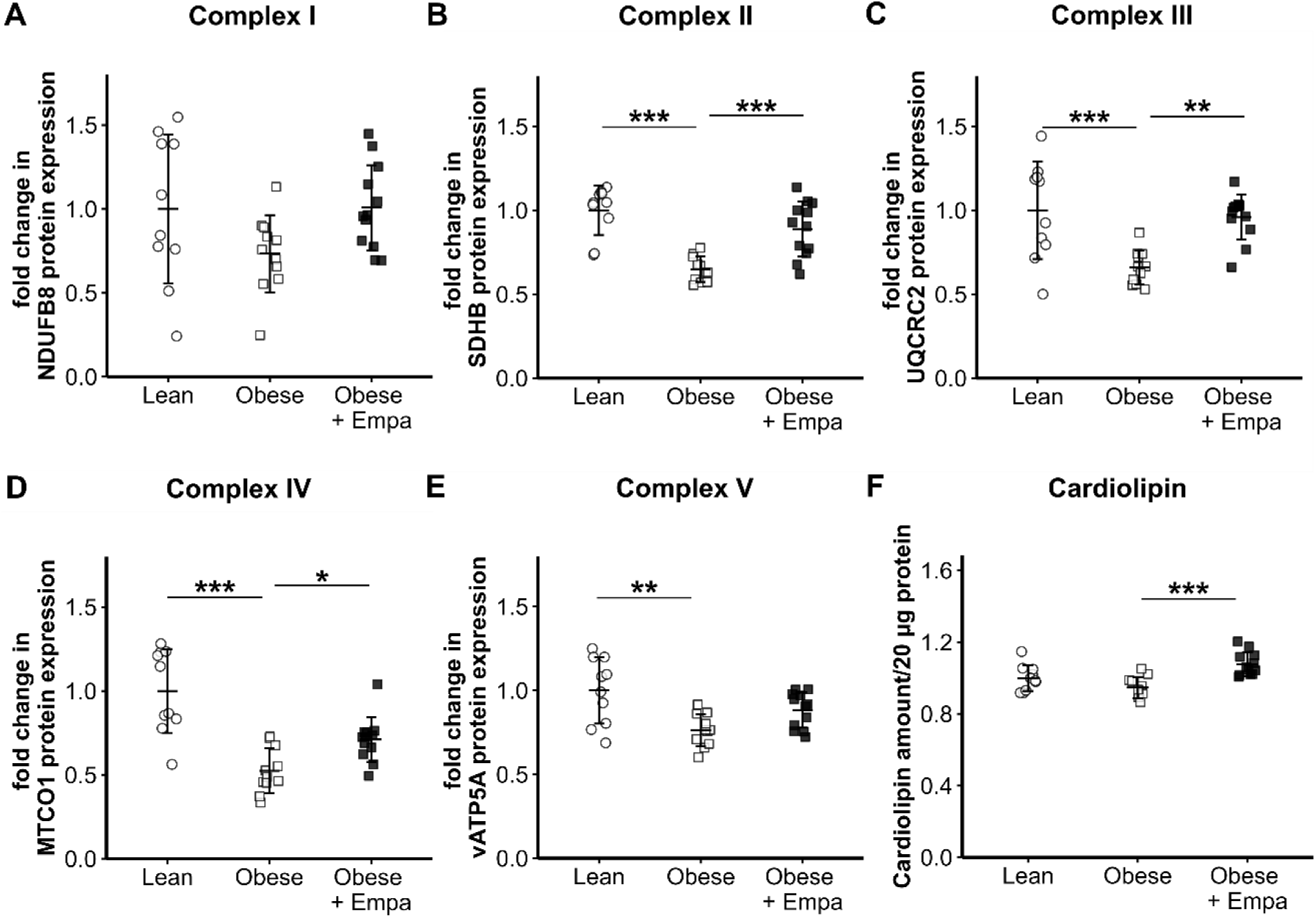
Protein expression of mitochondrial respiratory complex I-V. Protein expression of mitochondrial respiratory complexes I-V was evaluated in isolated kidney mitochondria of lean (white circles, n = 10), obese (white squares, n = 11) and empagliflozin-treated obese (black squaresn = 12) ZSF-1 rats. The protein expression was normalized to the house keeping gene porine and quantified as fold change to lean for A) complex I, the NADH dehydrogenase [ubiquinone] 1 beta subcomplex subunit 8 (NDUFB8) B) complex II, the succinate dehydrogenase [ubiquinone] iron-sulfur subunit (SDHB), C) complex III, the ubiquinol-cytochrome c reductase core protein 2 (UQCRC2), D) complex IV, the mitochondrial cytochrome c oxidase subunit I (MTCOI) and E) adenosine triphosphate 5A subunit (vATP5A). The corresponding representative western blot is shown in the supplement. F) Relative cardiolipin amount per 20 µg kidney mitochondria is shown as fold change versus lean group.

In addition, autophagy was evaluated by microtubule-associated protein 1A/1B light chain 3 I and II (LC3-I and LC3-II) expression. The obese animals had a significantly higher protein expression of LC3-I and LC3-II (Figure 6A, B), indicating the activation of autophagy. Treatment with empagliflozin resulted in a significant reduction in LC3-I and LC3-II protein expression. Protein expression of Fis-1, a marker of mitochondrial fusion and fission, was also assessed. The obese animals had significantly higher Fis-1 protein expression, which was significantly reduced after empagliflozin treatment (Figure 6C).

**Figure 6.**
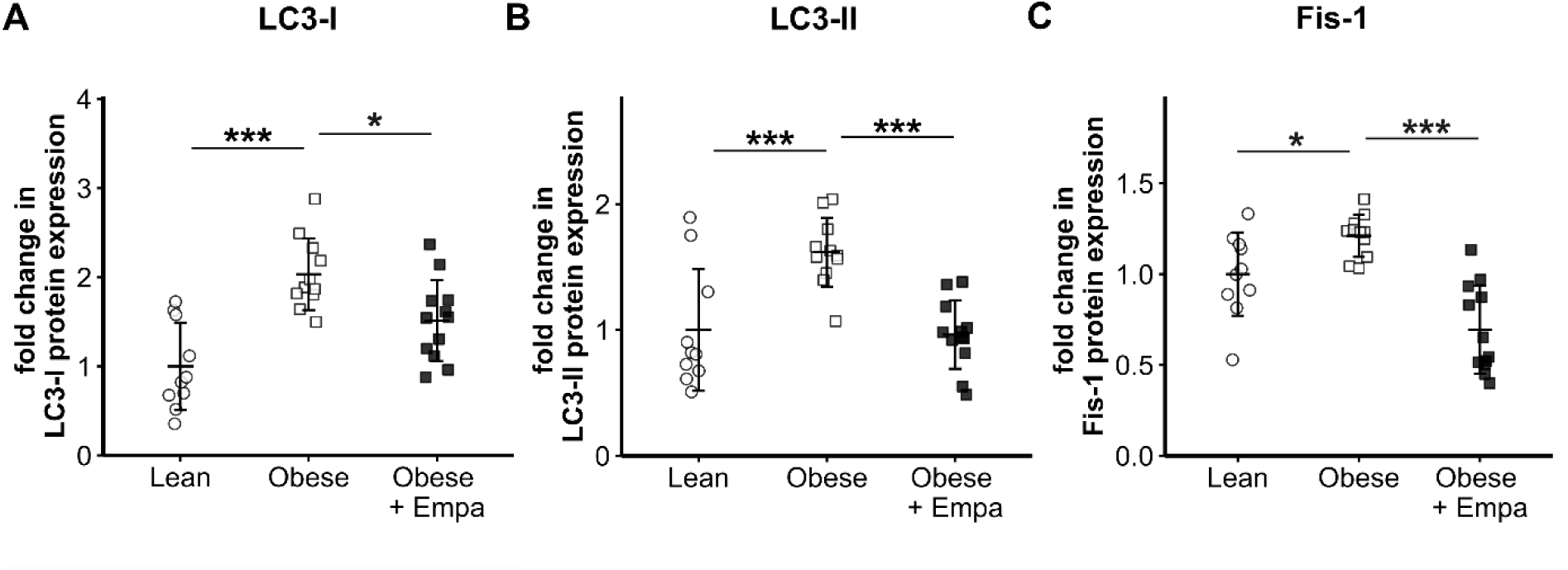
Protein expression of A) microtubule-associated protein light chain 3 (LC3) I, B) LC3-II and C) mitochondrial fission protein 1 (Fis-1). Protein expression was evaluated in isolated kidney mitochondria of lean (white circles, n = 10), obese (white squares, n = 11) and empagliflozin-treated obese (black squares, n = 12) ZSF-1 rats. The protein expression was normalized to the house keeping gene porine and quantified as fold change to lean. The corresponding representative western blots are shown in the supplement.

## Discussion

Metabolic dysregulation, characterized by conditions such as insulin resistance and obesity, can lead untreated to T2D. These comorbidities are encapsulated in the concept of metabolic syndrome, which causes vasoconstriction, increased renal sodium reabsorption, and elevated blood pressure. Consequently, this can result in CKD and cardiovascular disease [17,18]. The primary aim of the present study was to evaluate the effects of the SGLT2i empagliflozin in obese ZSF-1 rats. We specifically investigated how empagliflozin modulates diabetes-associated mitochondrial dysfunction and thereby may contribute to slow the progression of DN.

In T2D patients, early disease stages are characterized by glomerular hyperfiltration, followed by progressive GFR decline [19]. Similarly, obese ZSF-1 rats develop glomerular hyperfiltration as early as 10 weeks of age, followed by a age-dependent decline [20]. Consistent with previous findings [5,20], we did not detect significant differences in GFR at 24 weeks of age. However, by 32 weeks, obese rats exhibited a marked reduction in GFR, which was not attenuated by empagliflozin treatment, in line with previous studies [21]. Micropuncture studies by Vallon et al. demonstrated that SGLT2i reduce intraglomerular pressure and GFR in diabetic rats by restoring tubuloglomerular feedback primarily through afferent arteriolar vasoconstriction, but also with contribution of efferent arteriolar vasodilation [22]. In the present study, empagliflozin did not improve GFR after eight weeks of treatment, which might be the initial GFR decline (“GFR dip”) observed in clinical studies, which reflects acute hemodynamic changes thought to mediate long-term renoprotective effects [23]. In contrast, long-term treatment with ACE inhibitors for 24 weeks, targeting efferent arteriolar constriction, effectively preserved renal function and slowed disease progression in obese rats [5]. These findings suggest that in the obese ZSF-1 model, pathophysiological GFR changes might be predominantly driven by alterations in the efferent arteriole or are hidden behind the “GFR dip”, which may explain the limited effect of empagliflozin on GFR. Notably, despite the unchanged GFR, overall improvements of empagliflozin are present. Total proteinuria was significantly reduced, suggesting a renoprotective effect that may be at least partially independent of changes in overall glomerular filtration.

Furthermore, nephrin expression was significantly reduced in obese rats despite an unchanged number of WT1-positive podocytes, indicating slit diaphragm dysfunction rather than podocyte loss. Empagliflozin treatment restored nephrin expression, suggesting partial recovery of podocyte integrity, an essential component of the glomerular filtration barrier and a key mechanism underlying albuminuria. However, previous studies in other diabetic disease models, including diabetic db/db mice and obese ZSF-1 rats similarly reported that empagliflozin did not reduce albuminuria [21,24,25].

The observed reduction in total proteinuria with empagliflozin is linked to a decrease in urinary α1-microglobulin (α1M), a protein reabsorbed in the proximal tubule via receptor-mediated endocytosis. Elevated urinary α1M levels indicate impaired tubular reabsorptive function [26]. Its reduction signifies attenuation of tubular proteinuria by empagliflozin. This improvement in tubular protein handling aligns with the observed enhancement of tubular integrity, as demonstrated by reduced tubular cast formation, a hallmark of progressive kidney disease resulting from precipitation of filtered proteins within the tubular lumen [27–29]. These beneficial effects may be attributed to enhanced tubular reabsorptive capacity and reduced tubular workload, potentially secondary to reduced glucose reabsorption which may decrease tubular cast formation [12]. Despite its renoprotective effects, picrosirius red staining revealed persistent tubulointerstitial fibrosis following empagliflozin treatment. Collagen IV deposition was elevated in obese animals, but its progression was attenuated with empagliflozin. In contrast, another study in obese rats reported that empagliflozin promoted fibrosis via increased fibronectin accumulation without affecting collagen III deposition [21]. Thus, these divergent results may implicate that empagliflozin leads to a differential regulation of interstitial matrix expansion and glomerular basement membrane associated changes.

Reduction of metabolic stress in tubular cells is crucial for maintaining mitochondrial health. Mitochondrial ROS production is a major driver of tubular injury and disease progression and can induce uncoupling of mitochondrial respiration, thereby reducing ATP availability [13,30]. Previous studies have demonstrated impaired activity of complexes I, II, and IV, along with reduced protein abundance of complexes I and IV, in the context of T2D and obesity. In saponin-skinned myocardial and skeletal muscle fibers of obese ZSF-1 rats, empagliflozin improved mitochondrial function as well [31–33].

In our study, metabolic syndrome induced mitochondrial abnormalities, reflected by impaired respiratory capacity and altered OXPHOS protein expression. Although complex I respiration did not differ between lean and obese rats, empagliflozin significantly increased complex I state 3 (ADP-stimulated) respiration. As complex I represents the main entry point into the OXPHOS system and is essential for ATP production, its dysfunction contributes to renal injury [34] and has been linked to diabetes [35]. Improved complex I respiration was accompanied by enhanced substrate oxidation, more efficient ADP-to-ATP phosphorylation, and reduced proton leak [36]. Additionally, complex IV respiration and protein expression were significantly reduced in obese rats. Given that complex IV is required for complex I assembly and stability, its impairment may further compromise mitochondrial function [37]. Empagliflozin restored complex IV respiration and protein levels, suggesting improved OXPHOS integrity and overall mitochondrial stability. In contrast, ATP synthase protein expression was not restored by empagliflozin, which may indicate that metabolic injury had already caused persistent impairment of ATP-generating capacity. Sustained ATP deficiency can compromise Na⁺/K⁺-ATPase activity in the proximal tubule, impairing intracellular sodium homeostasis and contributing to hypertension and cardiorenal complications [38,39].

In addition, kidneys of obese ZSF-1 rats exhibited a pronounced shift toward fatty acid oxidation, which may overload mitochondria with β-oxidation derived intermediates, reduce metabolic flexibility and increase ROS production through elevated NADH and FADH₂ [40–42]. Empagliflozin shifts the whole-body substrate utilization toward fatty acids, reduces fat mass and lowers systemic burden [43]. This may relieve mitochondrial overload in the kidney, improve metabolic flexibility, reduce ROS generation, and support better mitochondrial function and tubular health.

Cardiolipin, a phospholipid of the inner mitochondrial membrane, is crucial for stabilizing electron transport chain complexes and supporting ATP production. [44,45]. Diabetes is known to induce cardiolipin remodeling and mitochondrial damage [46]. Empagliflozin treatment increased renal cardiolipin levels, accompanied by improved complex IV respiration and enhanced expression of respiratory complexes and mitochondrial quality control markers, indicating better mitochondrial function and dynamics. These effects may help maintain tubular integrity. Consistently, the cardiolipin-targeting antioxidant SS-31 has been shown to attenuate high-fat diet–induced glomerulopathy and proximal tubular injury by reducing mitochondrial damage and lipid accumulation [47]. Similar mitochondrial alterations were observed in the myocardial tissue of obese ZSF-1 rats, including reduced cardiolipin content and impaired respiratory capacity, which were attenuated by empagliflozin through stabilization of the respiratory chain [31].

In conclusion, and in contrast to the recently published paper by Hummelgaard et al., we observed renoprotective effects of empagliflozin in obese ZSF-1 rats. Despite the absence of GFR stabilization seen in other disease models and humans, we performed a much more detailed analysis of the morphological changes in the obese rats. Empagliflozin improved glomerular and tubular features, including podocyte function, tubular reabsorption and cast formation and mitochondrial health and function.

## Material & Methods

### Experimental Setup

All animal experiments were performed in accordance with the National Institutes of Health (NIH) *Guide for the Care and Use of Laboratory Animals* (NIH Pub. No. 85-23, Revised 1996) and the Federal Law on the Use of Experimental Animals in Germany and were approved by the local authorities (TVV 34/2020).

Female ZSF-1 obese rats and the corresponding lean controls were purchased from Charles River Laboratories, USA. Obese ZSF-1 rats were randomized at 24 weeks of age into a placebo-treated control group (Obese) and an empagliflozin-treated group (Obese + Empa) (8 weeks; 30 mg/kg bodyweight/day). Obese rats received standard diet and and were administered empagliflozin via chow. Lean, served as healthy controls and obese controls received standard chow and drinking water. At the age of 24 and 32 weeks, transdermal measurement of the glomerular filtration rate (GFR) was performed as previously described [10]. In brief, a fluorescence sensor was attached to the skin of conscious rats, fluorescein-isothiocyanate conjugated sinistrin was administered via the tail vein (40 mg/mL, 3 µL/g bodyweight) (MediBeacon, Germany) and the decrease in fluorescence intensity was measured over two hours by the sensors.

### Tissue Collection

At the age of 32 weeks, rats were anesthetized by intraperitoneal injection of 105 mg/kg bodyweight ketamine/7 mg/kg bodyweight xylazine. After opening the abdominal and thoracic cavities, the lower vena cava was punctured for blood collection. Kidneys were harvested and processed for further analysis. Blood samples were centrifuged for 15 min at 2,500 x *g*. Urine was collected by bladder puncture. Serum and urine were stored at -20°C and analyzed at the Institute for Clinical Chemistry and Laboratory Medicine at the University Hospital Carl Gustav Carus (Dresden, Germany) using standard laboratory methods.

### Renal Morphology

Kidney biopsies were fixed in zinc fixative over night at 4°C and were further processed as previously described [48]. Paraffin-embedded kidneys were cut into 2 µm thick sections, deparaffinized, rehydrated and stained with periodic acid–Schiff stain (PAS) or picrosirius red (PS) and hematoxylin as counter stain.

PAS-stained sections were automatically evaluated for glomerular size and PAS-positive glomerular area as previously described [49]. And manually scored for tubular damage using a tubular injury score (Supplement Table S3) [50]. For each animal 25 images of the whole kidney section were blinded scored for tubular injury. Tubular protein cast formation were counted in cortical and medullary region seperately, as a marker for tissue damage and progressive kidney disease and normalized the respective area.

Immunofluorescence staining was performed as previously described [51] using primary and secondary antibodies listed in Supplement Table S5 and counterstained with DAPI. Sections were scanned using the AxioScanZ.1 (Zeiss, Germany) at the Light Microscopy Facility of the CMCB of Technical University Dresden.

### Kidney Mitochondrial Respiration

For isolation of kidney mitochondria, cortical kidney tissue was homogenizing in cold homogenizing buffer using a teflon-homogenizer. Afterwards a series of centrifugation steps were performed (1. Cell homogenate at 500 x *g*, 10 min, 2. Supernatant at 10,000 x *g*, 10 min, 3. Resuspended pellet at 500 x *g*, 10 min, 4. Supernatant at 6,000 x *g*, 10 min, 5. Resuspended pellet at 500 x *g*, 10 min, 6. Supernatant at 5,000 x *g*, 10 min). All centrifugation steps were performed at 4°C. The final pellet was resuspended in 200 µL respiration medium. Composition of the buffers are listed in Supplement Table S2

The respiratory rates of renal cortical mitochondria were determined using a Clark electrode (Strathkelvin Instruments Limited, UK) as previously described [32]. The freshly isolated mitochondria were placed in the oxygraphic cell containing the respiration medium at 25°C with continuous stirring. The respiratory rates of the total mitochondrial population were measured twice by the sequential addition of specific substrates to evaluate the function of the different complexes of the OXPHOS system, as described in Supplementary Table S3. The respiratory control ratio (RCR) for complex I was calculated as the ratio of state 3 to state 2 respiration.

### Western Blot Analysis

For western blot analysis, a total of 4 µg isolated kidney mitochondria were used. Protein concentration was determined using a Pierce^TM^ BCA Protein Assay Kit (ThermoFisher Scientific, Germany). Protein samples were separated by SDS-polyacrylamide gel electrophoresis. Proteins were then transferred to a polyvinylidene fluoride (PVDF) membrane, incubated overnight at 4°C, washed and then incubated with secondary antibodies for 2 hours at room. temperature Antibodies are listed in Supplement Table S4. The proteins were detected using ECL Western Blotting Substrate (Promega Corporation, USA) and visualized using Fusion FX Imager. Protein quantification was performed using Bio1D (Vilber Lourmat Deutschland GmbH, Germany). Measurements were normalized to the house keeping gene porine, a protein that is expressed in the outer mitochondrial membrane

### RNA Isolation

Total RNA was isolated from kidney samples using TRIzol™ reagent (Invitrogen Inc., USA). Kidney tissue was homogenized in 1 mL of TRIzol™ using a bead homogenizer, followed chloroform extraction (0.2 ml) and centrifugation (30 min, 4°C, 12,000 x *g*). The aqueous phase was transferred to a new tube, and 0.5 ml 2-propanol was added, incubated on ice for and centrifuged again (10 min, 4°C, 12,000 x *g*). The RNA pellets were washed with ice-cold 75% ethanol, centrifuged (10 min, 4°C, 12,000 x *g*), air dried for 30 min and dissolved in 60 µL RNase-free water at 55°C for 15 min. RNA concentration was determined using a NanoDrop™ 2000c (Thermo Fisher Scientific Inc., Germany) and samples were stored at - 80°C until further analysis.

### Reverse Transcription and quantitative Real-Time PCR Analysis

One µg total RNA was used for cDNA synthesis using the SensiFAST™ cDNA Synthesis Kit (Meridian Bioscience Inc., USA) according to manufacturer’s instructions. The SensiFAST™ SYBR® No-ROX Kit (Meridian Bioscience Inc., USA) was used for quantitative real-time PCR analysis (T-Optical, Jena Analytics, Germany). A QuantiTect Primer Assay Kit (Qiagen GmbH, Germany QT00180852) specific for rat *Slc5a2* was used.

### α1-Microglobulin ELISA

The α1-Microglobulin ELISA Kit (antibodies.com, A78985) was performed according to the manufacturer’s instructions.

### Cardiolipin Assay

The cardiolipin assay (abcam, ab241036) was performed using 20 µg isolated kidney mitochondria according to the manufacturer’s instructions.

### Statistics

The data were visualized and analyzed using R (version 4.0.2, [52]) with RStudio (version 1.2.5033, [53]). A one-way ANOVA was performed to compare multiple groups, followed by the Bonferroni multiple comparison test. P values <0.05 were considered statistically significant.

## Acknowledgments

We highly acknowledge the technical assistance of Anika Wirth, Andrea Angermann, Katja Maiwald and Maria Schuster.

## Authors contributions

AS, FG, VA and CH designed the study. HW, AS, AM, ASt, JM, MS and FG performed experiments and HW did the data analysis. HW wrote the manuscript. All authors have approved the manuscript. VA and AS contributed equally.

## Data Availability Statement

The data that support the findings of this study are available from the corresponding author upon reasonable request.

## Ethics Statement

Not applicable.

