## Supplemental material for "Empagliflozin preserves mitochondrial function and reduces tubular injury in obese type 2 diabetic ZSF-1 rats"

### Supplement Figures

**Table S1 Tubular injury score.** Score for evaluation of tubular injury in ZSF-1 rats.

| **Score** | **Damage** |
|---|---|
| **0** | No tubular damage |
| **1** | Loss of brush border integrity in less than 25% of tubular cells and integrity of the basal membrane |
| **2** | Loss of brush border integrity in more than 25% of tubular cells and thickening of the basal membrane |
| **3** | Additional inflammation, cast formation |

**Table S2 Buffer used for kidney mitochondrial respiration.**

| Homogenizing Buffer pH 7.4 | Concentration |
|---|---|
| Sucrose | 250 mmol/L |
| Tris | 10 mmol/L |
| EGTA, | 500 mmol/L |

| Respiration medium, pH 7.1 | |
|---|---|
| EGTA | 0.5 mmol/L |
| $MgCl_2$ | 3 mmol/L |
| Taurine | 20 mmol/L |
| $KH_2PO_4$ | 10 mmol/L |
| HEPES | 20 mmol/L |
| sucrose | 110 mmol/L |
| potassium lactobionate solution | 60 mmol/L |
| BSA | 1 g/L |

**Table S3 Substrates used for the measurement of mitochondrial respiration.** Mitochondrial respiration was measured in two different setups and the listed substrates were added sequentially.

| | |
|---|---|
| **Setup number 1** | |
| | **Substrate (Concentration)** |
| **1** | glutamate/malate (10 mmol/L / 2 mmol/L) |
| **2** | Adenosine diphosphate (2 mmol/L) |
| **3** | Antimycin A (2.5 µmol/L) |
| **4** | ascorbate/N,N,N’,N’-tetramethyl-p-phenylenediamine dichloride (2 mmol/L / 0.5 mmol/L) |
| **5** | Sodium azide (4 mol/L) |
| **Setup number 2** | |
| | **Substrate (Concentration)** |
| **1** | glutamate/malate (10 mmol/L / 2 mmol/L) |
| **2** | Adenosine diphosphate (2 mmol/L) |
| **3** | Octanoylcarnithine (100 µmol/L) |
| **4** | Succinate (10 mmol/L) |

**Table S4 Antibodies used for western blot analysis.**

| Primary antibodies | Dilution | Distributor |
|---|---|---|
| Total OXPHOS Rodent WB Antibody Cocktail | 1:250 | Abcam |
| Fis1 | 1:1,000 | Proteintech |
| LC3B | 1:1,000 | Sigma |
| Porine | 1:1,000 | Abcam |
| **Secondary antibodies** | | |
| anti-mouse IgG HRP | 1:5,000 | Abcam |
| anti-rabbit IgG peroxidase | 1:10,000 | Sigma |

**Table S5 Antibodies used for immunofloureszenz stainings.**

| Primary antibodies | Dilution | Distributor |
|---|---|---|
| Nephrin | 1:100 | Progen (GP-N2) |
| Collagen IV | 1:200 | Southern Biotechnology Ass. (1340-01) |
| WT1 | 1:200 | abcam (ab29) |
| Cubilin | 1:100 | Santa Cruz Technology (sc-20609) |
| **Secondary antibodies** | | |
| anti-guinea pig 647 | 1:500 | Jackson Immuno (706-605-148) |
| anti-goat 555 | 1:500 | Invitrogen (A-21432) |
| anti-rabbit 647 | 1:500 | abcam (ab150075) |

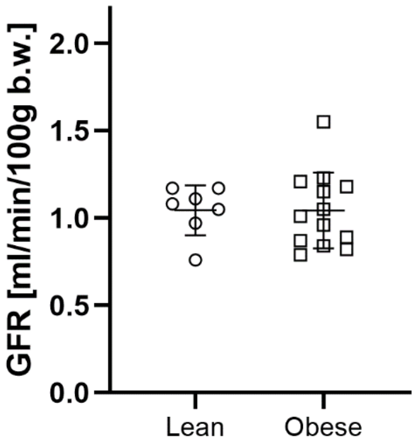


**Figure S 1 Glomerular Filtration Rate (GFR) at 24 weeks of age in lean and obese rats.**

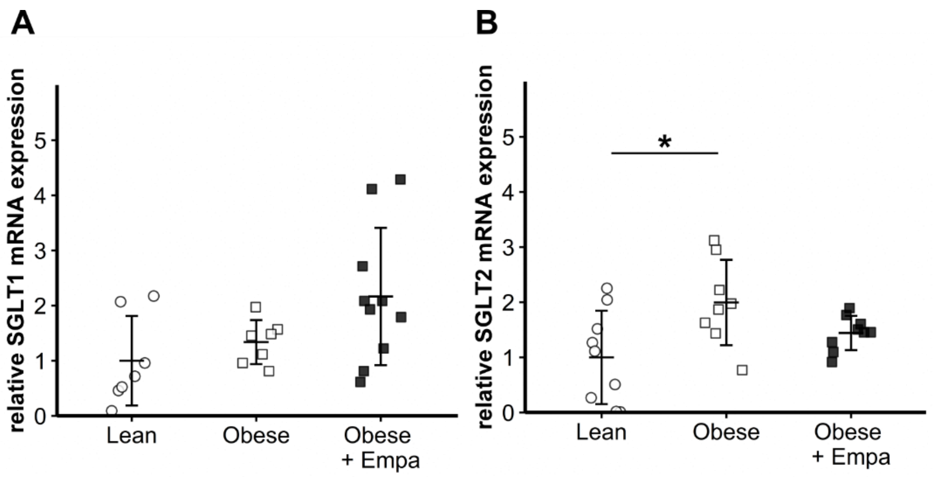


**Figure S 2 Renal SGLT1 and SGLT2 mRNA expression.** A) Relative SGLT1 mRNA expression and B) relative SGLT2 mRNA expression in lean (white circles), obese (white squares) and empagliflozin-treated obese (black squares) ZSF-1 rats. mRNA expression levels were normalized to the hypoxanthine guanine phosphoribosyl transferase (*Hprt*). Relative mRNA expression is shown as fold change versus lean group. Number of animals ≥ 12 in each group.

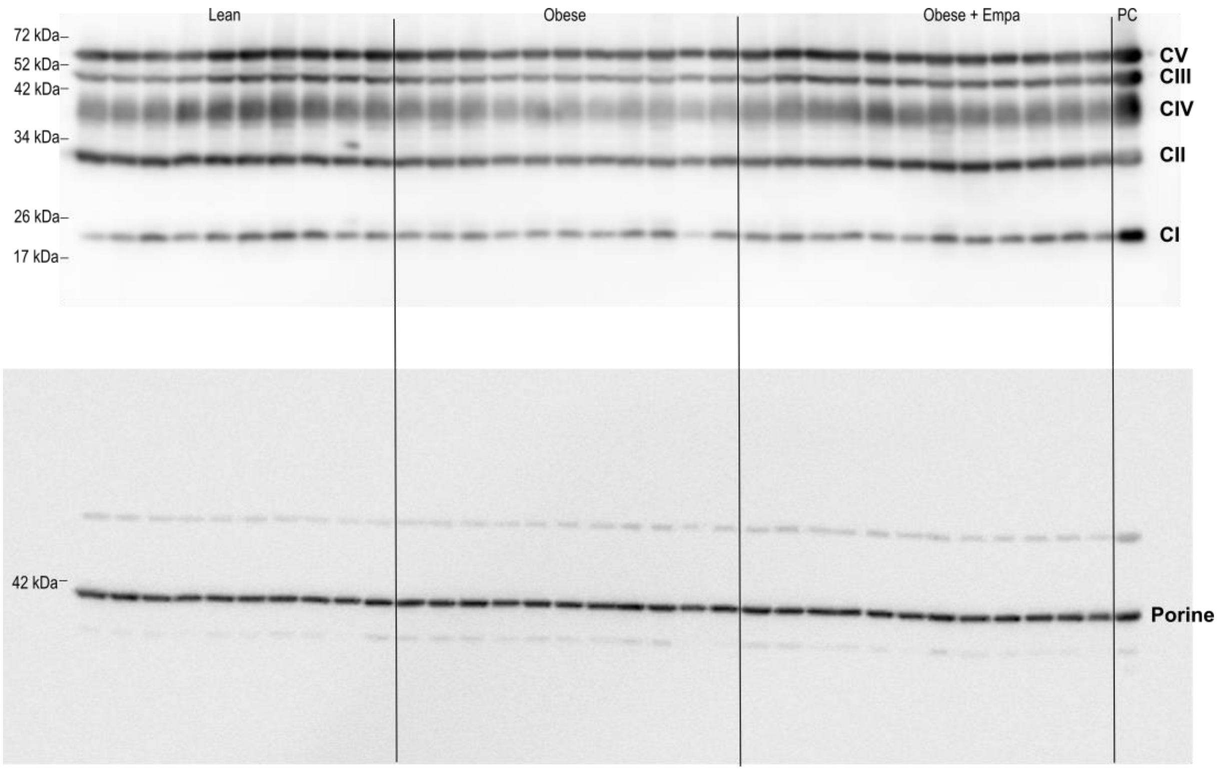


**Figure S 3 Uncropped western blots of mitochondrial respiratory complex I-V and the house keeping gene porine.** CI-Complex I, CII-Complex II, CIII- Complex III, CIV-Complex IV, CV-Complex V, PC-Positive control (delivered with the antibody).

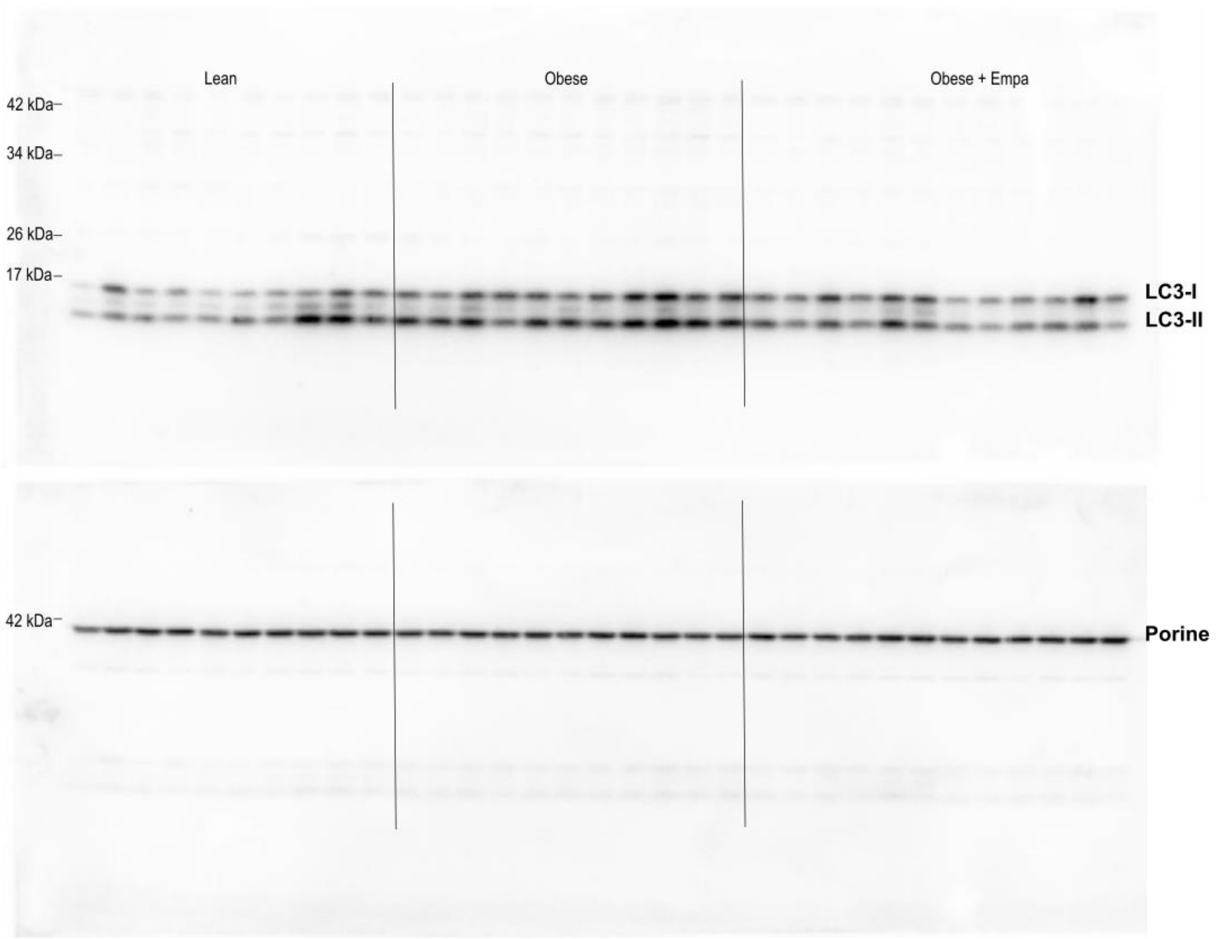


**Figure S 4 Uncropped western blot of LC3-I and LC3-II and the house keeping gene porine.**

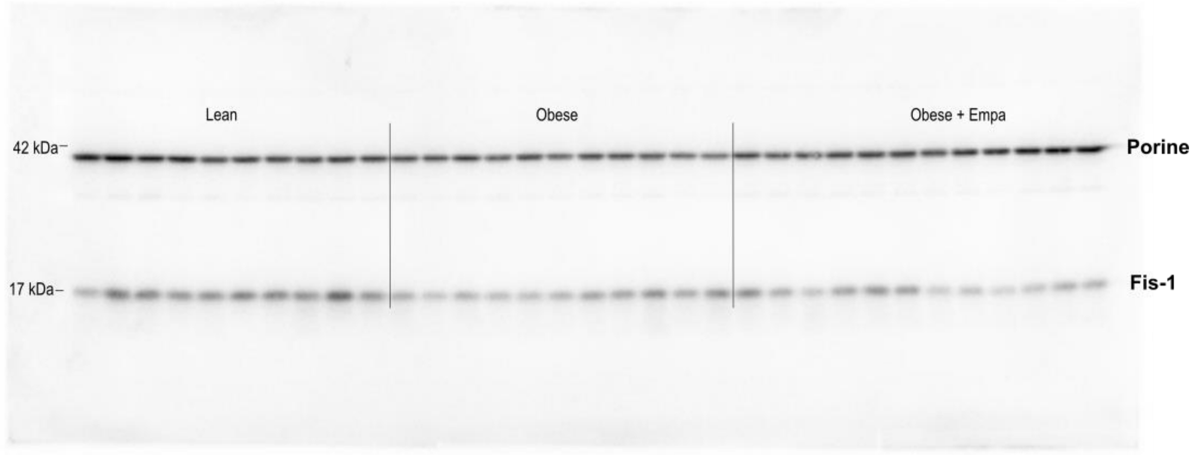


**Figure S 5 Uncropped western blot of Fis-1 and the house keeping gene porine.**
